# Intrinsic neural timescales related to sensory processing: Evidence from abnormal behavioural states

**DOI:** 10.1101/2020.07.30.229161

**Authors:** Federico Zilio, Javier Gomez-Pilar, Shumei Cao, Jun Zhang, Di Zang, Zengxin Qi, Jiaxing Tan, Tanigawa Hiromi, Xuehai Wu, Stuart Fogel, Zirui Huang, Matthias R. Hohmann, Tatiana Fomina, Matthis Synofzik, Moritz Grosse-Wentrup, Adrian M. Owen, Georg Northoff

## Abstract

The brain exhibits a complex temporal structure which translates into a hierarchy of distinct neural timescales. An open question is how these intrinsic timescales are related to sensory or motor information processing and whether these dynamics have common patterns in different behavioural states. We address these questions by investigating the brain’s intrinsic timescales in healthy controls, motor (amyotrophic lateral sclerosis, locked-in syndrome), sensory (anaesthesia, unresponsive wakefulness syndrome), and progressive reduction of sensory processing (from awake states over N1, N2, N3). We employed a combination of measures from EEG resting-state data: auto-correlation window (ACW), power spectral density (PSD), and power-law exponent (PLE). Prolonged neural timescales accompanied by a shift towards slower frequencies were observed in the conditions with sensory deficits, but not in conditions with motor deficits. Our results establish that the spontaneous activity’s intrinsic neural timescale is related to specifically sensory rather than motor information processing in the healthy brain.

**Highlights:**

- EEG resting-state shows a hierarchy of intrinsic neural timescales.
- Sensory deficits as in disorders of consciousness lead to prolonged intrinsic neuraltimescales.
- Clinical conditions with motor deficits do not show changes in intrinsic neural timescales.20

**Graphical Abstract:** 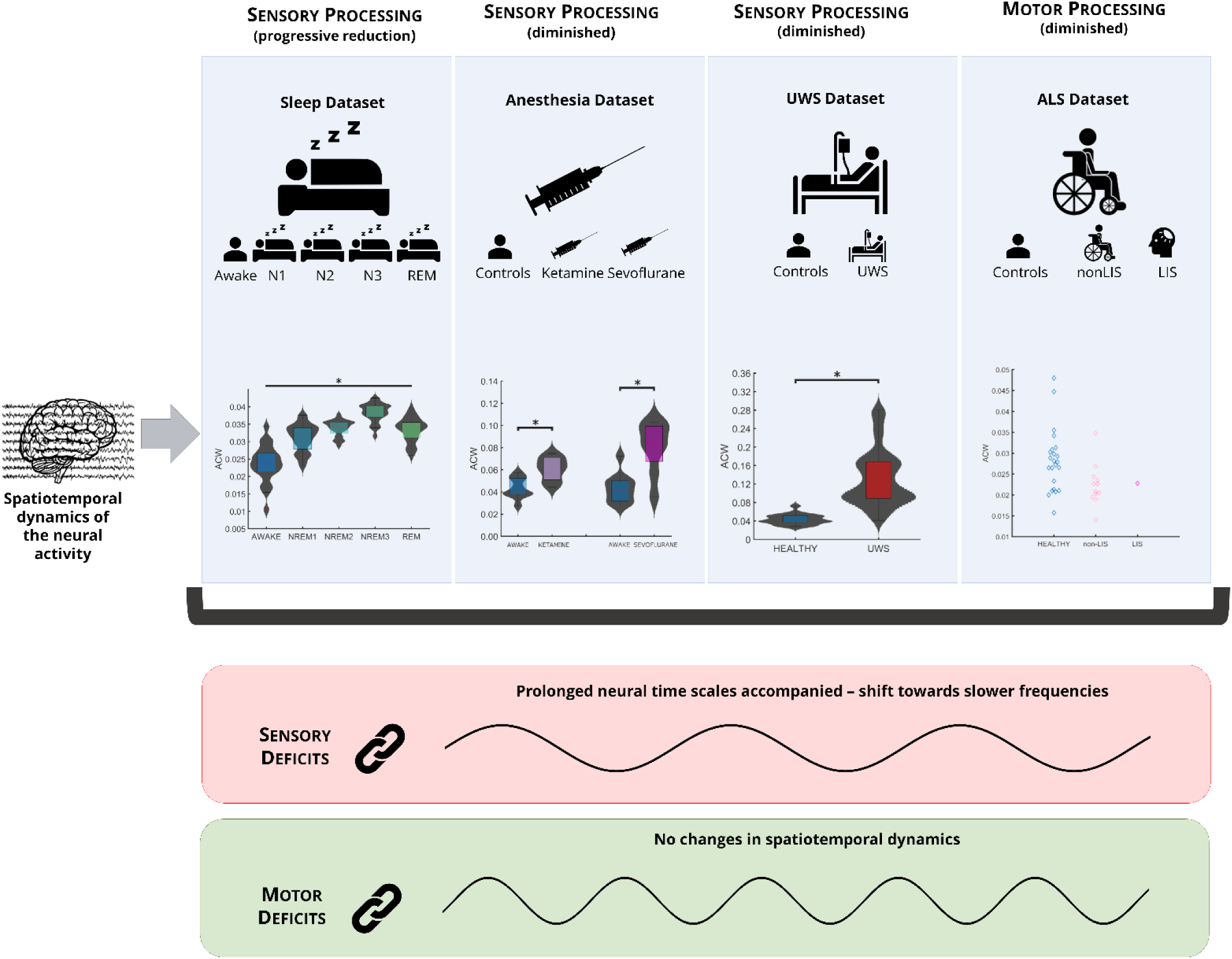

## Introduction

### Intrinsic neural timescale

The spatiotemporal dynamics of a neural system shape information processing. Information is processed by the brain’s intricate temporal structure. In other words, much like a radio can receive messages by decoding the modulation of the amplitude (AM) or frequency (FM) of radio signals, the language that the brain uses to communicate with itself is encoded/structured in time. We consequently need to investigate the temporal-spatial dynamics of the brain in order to understand the fundamental processes of healthy and disordered states of consciousness. Different brain regions exhibit different “temporal receptive fields” (Cavanagh et al., 2016) or “temporal receptive windows” (Bernacchia et al., 2011; Chaudhuri et al., 2015; Chen et al., 2016, 2015; Cocchi et al., 2016; Demirtaş et al., 2019; Farzan et al., 2017; Gollo et al., 2017, 2015; Hasson et al., 2015; Honey et al., 2012; Huang et al., 2018a; Kiebel et al., 2008; Luppi et al., 2019; Mohr et al., 2016; Murray et al., 2014; Runyan et al., 2017; Stephens et al., 2013; Wasmuht et al., 2018; Watanabe et al., 2019; Wolff et al., 2019). These findings led to the assumption that different regions and networks in the brain exhibit their specific timescales as reflected in the concept of “intrinsic neural timescales” (Chaudhuri et al., 2015; Deco et al., 2019; Farzan et al., 2017; Gollo et al., 2017, 2015; Liégeois et al., 2019; Murray et al., 2014; Wasmuht et al., 2018).

The length of intrinsic neural timescales differs from one brain region to another. For example, the intrinsic neural timescales are shorter in sensory and motor regions while they seem to be longer in higher-order cortical regions (Murray et al., 2014; Ogawa and Komatsu, 2010; Stephens et al., 2013). In addition, brain regions that support temporal pooling and summation (Himberger et al., 2018) of sensory (Gauthier et al., 2012; Hasson et al., 2008; Lerner et al., 2011; Stephens et al., 2013; Yeshurun et al., 2017), motor, and cognitive information (Bernacchia et al., 2011; Farzan et al., 2017; Hasson et al., 2015; Murray et al., 2014) have unique temporal signatures. Thus, these distinct intrinsic timescales may provide a meaningful functional dissociation between brain areas. However, how these intrinsic neural timescales modulate and integrate information (e.g., sensory vs. motor) remains an open question in systems neuroscience. Addressing this question is the main aim of the current investigation.

Current evidence for a role of intrinsic neural timescales in information processing is largely indirect, stemming mainly from the use of temporal measures in pathological cases. For example, abnormal intrinsic timescales from resting-state fMRI in psychiatric disorders such as autism (Damiani et al., 2019; Watanabe et al., 2019) are accompanied by deficits in sensory processing and abnormal social behaviour. Yet another study, using resting-state fMRI, demonstrated abnormally long intrinsic timescales in conditions involving reduced or absent sensory behaviour, e.g., anaesthesia, unresponsive wakefulness state (UWS), minimally conscious state (MCS), or non-rapid eye movement (NREM) sleep stages (N1-3) (Huang et al., 2018a). Nevertheless, there is a need for research employing more direct measures of the brain’s intrinsic neural timescale probing their involvement in specific functions like sensory or motor functions. For example, measures that quantify the frequency characteristics of the periodic oscillations employed in the current study (e.g., power spectral density; PSD), arrhythmic scale-free (“1/f noise”) brain activity (e.g., power-law exponent; PLE), and the repeating patterns in a signal (e.g., autocorrelation window; ACW) are novel and potentially powerful ways to explore the temporal structure of neuronal communication at the systems level.

### Metrics of temporal dynamics

To investigate the intrinsic neural timescale of resting-state EEG, we calculated autocorrelation using a well-established measure, the ACW. The ACW measures repeating patterns in a signal, and enables us to test for the relationship, e.g., correlation in neural activity patterns at different points in time (Murray et al., 2014). The ACW has been applied at both the cellular (Bernacchia et al., 2011; Cavanagh et al., 2016; Murray et al., 2014) and systems levels (Huang et al., 2018b; Watanabe et al., 2019; Wolff et al., 2019). Therefore, the ACW can be regarded as a valid and direct measure of the intrinsic neural timescale. Moreover, it has recently been suggested that the ACW is related to slow frequencies (Honey et al., 2012). Thus, we also measured the frequency characteristics of the periodic oscillations in EEG using PSD and the arrhythmic scale-free (“1/f noise”) brain activity using PLE in our various groups (He, 2014; He et al., 2010; Huang et al., 2017a, 2016; Linkenkaer-Hansen et al., 2001; Palva and Palva, 2018). The additional measurements of the PSD and the PLE allowed us to link the ACW to power in different frequencies across normal and disordered states of consciousness. Exploring the relationship between the ACW and the frequency characteristics of the EEG will enable us to identify the supposed role of intrinsic timescale in temporal integration of sensory or motor stimuli (Florin and Baillet, 2015; Himberger et al., 2018).

### Processing of sensory vs. motor information

Unresponsive Wakefulness State (UWS) and Minimally Conscious State (MCS): The frequency characteristics of intrinsic neural timescales can be measured using the PSD (Chaudhuri et al., 2015; Murray et al., 2014; Rosanova et al., 2018; Wolff et al., 2019). Slowing in the PSD would reflect prolongation of the intrinsic neural timescales, and would be expected in cases of UWS, MCS, anaesthesia, and slow wave sleep (N3). Behaviourally, despite their differences, UWS and MCS share the loss of sensory function as seen by reduced sensory-evoked potentials (Banoub et al., 2003; Boisseau et al., 2002; Boly et al., 2008, 2004; Fischer et al., 2010; Nakano et al., 1995; Noguchi et al., 1995; Pistoia et al., 2016; Rosanova et al., 2018; Schiff et al., 2014; Sharon and Nir, 2018; Wang et al., 2003; Wijnen et al., 2014; Xu et al., 2012). In contrast, motor function (e.g., involuntary movements and motor-evoked potentials) in UWS and MCS remain intact, except in slow wave sleep (Bergmann et al., 2012). This suggests that abnormal prolongation of the intrinsic neural timescales will be associated with a deficit, or loss, of the capacity for sensory information processing, rather than for motor processing, although this possibility remains to be explored. Locked-in-Syndrome (LIS) and Amyotrophic Lateral Sclerosis (ALS): In contrast to UWS and MCS, locked-in-syndrome (LIS) and amyotrophic lateral sclerosis (ALS) present the opposite behavioural pattern. In the case of LIS and ALS, sensory function including somatosensory-evoked potentials and brain-stem auditory evoked potentials remain completely, or partially intact (Bassetti et al., 1994; Behr et al., 1991; Bensch et al., 2014; Facco et al., 1989; Gosseries et al., 2009; Hammond and Wilder, 1982; Landi et al., 1994; Soria et al., 1989; Virgile, 1984). However, motor-evoked potentials are lost in LIS and ALS (Bassetti et al., 1994; Facco et al., 1989; Kotchoubey and Lotze, 2013; Landi et al., 1994). The loss of motor function in LIS and ALS is due to the disruption of descending motor pathways, despite intact activation in cortical motor areas (Cincotta et al., 1999). Movement-related alterations of PSD in the beta band have also been reported in ALS (Bizovičar et al., 2014; Proudfoot et al., 2018, 2017). However, no studies have investigated intrinsic timescales in LIS and ALS (see though (Babiloni et al., 2010; Proudfoot et al., 2018, 2017) for observed power changes in alpha, beta and gamma bands).

If it is indeed the case that intrinsic timescales are related to sensory (rather than motor) processing (as in UWS and MSC), one would predict no changes in the intrinsic neural timescales in primarily motor conditions like LIS and ALS where sensory processing is intact. To address this, we compared the intrinsic neural timescales in primarily motor-deficient but sensory-preserved conditions (LIS, ALS) to those of primarily sensory-deficient but motor-preserved behavioural conditions (anaesthesia, MCS/UWS) and also conditions involving normal and healthy reduced motor activity and progressive alterations of sensory processing (e.g., slow wave sleep). This approach will enable us to determine whether the intrinsic neural timescale of the brain’s spontaneous activity is central for either sensory or motor information processing, or both.

### General and specific aims

The overarching aim of our study was, therefore, to use EEG resting-state to investigate the relationship between the intrinsic neural timescale of the brain’s spontaneous activity and sensory or motor information processing.

The first specific aim was to probe the ACW (as well as the PLE and the PSD) in the EEG resting-state of sensory-deficient but motor-preserved behavioural conditions comparing them with sensory-preserved healthy states. These conditions included anaesthesia, sleep, and UWS, where sensory information processing is reduced or lost in either naturally (sleep, UWS) and non-naturally (anaesthesia) occurring states. Previous findings report abnormal temporal dynamics with slowing of the PSD (and/or high PLE) in anaesthesia, sleep, and UWS (Akeju et al., 2016; Casali et al., 2013; Demertzi et al., 2019; Huang et al., 2018c, 2016, 2014; Tagliazucchi et al., 2016, 2013b). Based on these findings, we hypothesized that the ACW would be longer in sleep, anaesthesia, and UWS when compared to fully awake states in either the same participant (sleep, anaesthesia) or some healthy control group (UWS). This longer duration of the ACW would suggest that neural activities at more distant time points strongly correlate, and thus strongly resemble each other. Together with the supposed shift towards slower frequencies in the PSD and the PLE, this similarity across distant time points increases the processing capacity for temporal integration of temporally distant sensory information. This, in turn, reduces the temporal precision of specific sensory information at specific discrete points in time with the subsequent loss of perception of specific objects or events (Himberger et al., 2018).

The second specific aim was to probe the ACW (as well as the PLE and the PSD) in the EEG resting-state of motor-deficient but sensory-preserved behavioural conditions comparing them with motor-preserved healthy states. This was done by comparing primarily motor-deficient conditions like ALS and LIS with healthy controls. Given the evidence that the ACW may be central in sensory rather than motor information processing (see above), we hypothesized that there would be no differences in the ACW duration (nor in the PLE and the PSD) in ALS and LIS when compared to healthy control participants. In order to test for the hypothesis of no difference, we employed statistical tests of equivalence for non-inferiority. We here assessed the temporal structure of the intrinsic neural timescale of the brain’s spontaneous activity using the ACW, PLE and PSD in different behavioural conditions and states of consciousness. This allowed us to address the question of whether the intrinsic timescales of the brain’s spontaneous activity support the processing of specific information, that is, sensory or motor information. We used a comparative approach in healthy fully awake brain state and sensory-and motor-compromised states. In this way, we hoped to reveal how the hierarchical structure of intrinsic timescales support temporal integration (or segregation) of sensory rather than motor information processing across normal and abnormal states of consciousness. In addition to revealing the functional role of intrinsic timescales, this also provides insight into the neural correlates that support conscious arousal and awareness under healthy conditions.

## Material and methods

### Participants

Following the general aim of investigating intrinsic EEG timescales in different behavioural conditions, four datasets were analysed: (1) sleep dataset, (2) anaesthesia dataset, (3) UWS dataset, and (4) amyotrophic lateral sclerosis (ALS) dataset. A description of each follows and are presented in Table 1.

**Table 1.**
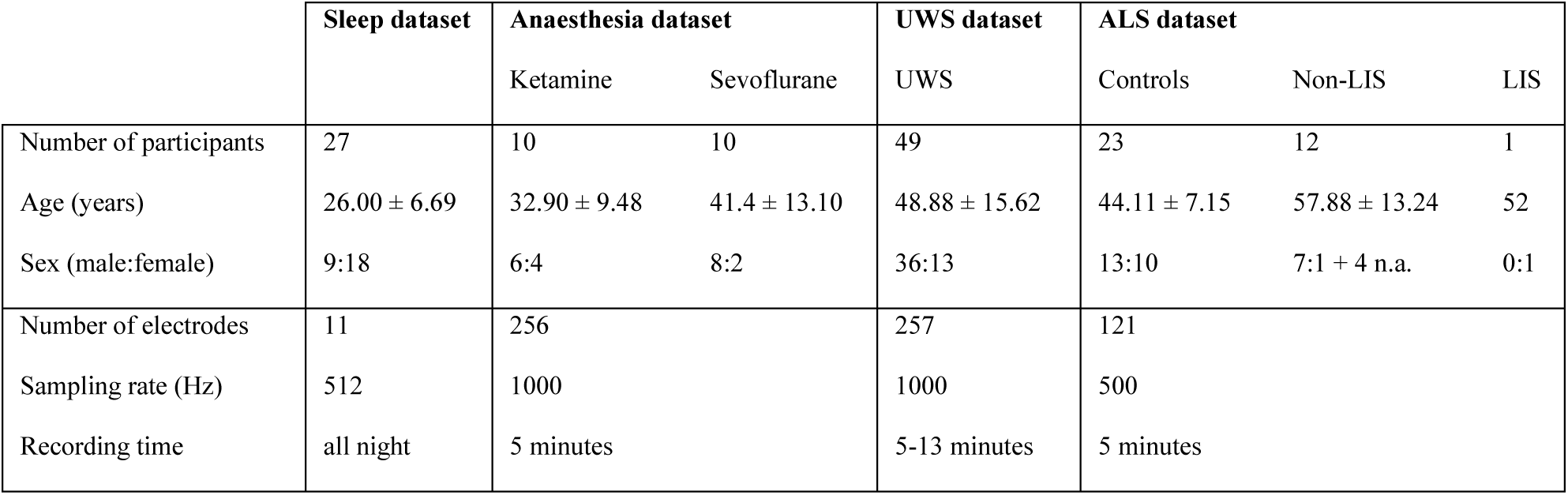
Summary of the main characteristics of each dataset.

|  | Sleep dataset | Anaesthesia dataset |  | UWS dataset | ALS dataset |  |  |
| --- | --- | --- | --- | --- | --- | --- | --- |
|  |  | Ketamine | Sevoflurane |  | Controls | Non-LIS | LIS |
| Number of participants | 27 | 10 | 10 | 49 | 23 | 12 | 1 |
| Age (years) | $26.00 \pm 6.69$ | $32.90 \pm 9.48$ | $41.4 \pm 13.10$ | $48.88 \pm 15.62$ | $44.11 \pm 7.15$ | $57.88 \pm 13.24$ | 52 |
| Sex (male:female) | 9:18 | 6:4 | 8:2 | 36:13 | 13:10 | 7:1 + 4 n.a. | 0:1 |
| Number of electrodes | 11 | 256 |  | 257 | 121 |  |  |
| Sampling rate (Hz) | 512 | 1000 |  | 1000 | 500 |  |  |
| Recording time | all night | 5 minutes |  | 5-13 minutes | 5 minutes |  |  |

### Sleep dataset

Twenty-seven, healthy adults (age =26.00 ± 6.69 years, 18 women) were included in this study. All participants reported normal sleep patterns, and were free from signs of sleep disorders, according to standard guidelines (AASM, 2014), assessed from an overnight polysomnographic (PSG) screening night. Participants performed a complete PSG using the Embla Titanium (Natus, San Carlos, CA) PSG system. EEG, EOG and EMG signals were acquired with impedances < 5 KΩ, at a sampling rate of 512 Hz, referenced to FPz. EEG was acquired using 11 gold-plated electrodes placed according to the conventional 10-20 system. The EEG signals were re-referenced offline to the average of the mastoid derivations for sleep stage scoring. Sleep stages (wake before sleep, N1, N2, N3, REM) were marked using RemLogic analysis software (Natus) following the standard criteria (Iber et al., 2007).For a further description of the dataset, see (Fang et al., 2017a).

### Anaesthesia dataset

For the anaesthesia dataset, the effects of two different general anaesthetics, i.e. ketamine and sevoflurane were evaluated. To assess the ketamine effect, resting-state EEG recordings (Ges300, EGI, USA) in ten right-handed surgical patients aged between 20-60 years (32.90 ± 9.48 years, 4 women), American Society of Anesthesiologists (ASA) physical status class I - II, were collected in awake (5 min eyes-closed) condition using an electrode cap (HydroCel 130) of 256 electrodes following 10-20 international system. Then, ketamine was given to the same 10 participants. Specifically, 1 mg/kg diluted ketamine in 10 ml of 0.9% normal saline was infused over 2 min, until OAA/S (Observer’s Assessment of Alertness/Sedation) scale was 1. Then, ultrashort-acting opioid remifentanil 1μg/kg and neuromuscular relaxant rocuronium 0.6mg/kg were given for endotracheal intubation. After anaesthetic induction, diluted ketamine was infused again over 20 min (1 mg/kg/h). EEG data (5 minutes) were acquired again from 15 min after the loss of consciousness. In order to avoid external noise interference, all participants were placed earplugs in both ears. During the EEG acquisition at a sampling rate of 1000 Hz, electrode impedance was kept under 5 KΩ. All channels were referenced online to Cz. Similarly, ten different participants (age = 41.4 ± 13.10 years, 2 women) followed the same protocol but under sevoflurane anaesthesia. In this case, 8% sevoflurane was initially administered in 6L/min 100% oxygen and when OAA/S score was 1, remifentanil 1μg/kg and rocuronium 0.6mg/kg was given for endotracheal intubation. After anaesthetic induction, the end-tidal concentration of sevoflurane was kept at 1.3 MAC (2.6%). EEG data (5 minutes) were acquired from 15 min after the loss of consciousness. Equipment and EEG acquisition procedures were the same as the ones followed under the effects of ketamine. During the study period, electrocardiogram, non-invasive blood pressure and pulse oximetry were monitored in these non-premedicated participants (see Table 2 for further details).

**Table 2.** The clinical data before and after anaesthesia in two anaesthesia groups.

| Parameter | Awake state | Anaesthetic state | P-value |
| --- | --- | --- | --- |
| **Sevoflurane anaesthesia** |  |  |  |
| HR (beats/min) | 69.5±7.6 | 67.8±7.5 | 0.37 |
| SBP (mmHg) | 125.2±13.7 | 101.2±23.7 | <0.01 |
| DBP (mmHg) | 71.2±9.6 | 56.3±17.9 | <0.01 |
| RR (times/min) | 13.2±1.6 | 10.7±1.0 | 0.01 |
| SpO<sub>2</sub> (%) | 98.5±1.4 | 99.0±0.6 | 0.08 |
| PaO<sub>2</sub> (mmHg) | 105.5±17.4 | 451.5±155.8 | <0.01 |
| PaCO<sub>2</sub> (mmHg) | 38.9±3.5 | 39.1±3.7 | 0.28 |
| PH | 7.43±0.05 | 7.42±0.01 | 0.55 |
| **Ketamine anaesthesia** |  |  |  |
| HR (beats/min) | 75.2±13.0 | 86.3±14.0 | 0.02 |
| SBP (mmHg) | 137.5±18.3 | 151.5±18.0 | <0.01 |
| DBP (mmHg) | 73.2±15.2 | 84.9±8.3 | 0.01 |
| RR (times/min) | 13.4±1.6 | 11.4±1.3 | 0.04 |
| SpO<sub>2</sub> (%) | 98.1±1.4 | 99.2±0.6 | 0.08 |
| PaO<sub>2</sub> (mmHg) | 105.5±12.1 | 451.5±150.8 | <0.01 |
| PaCO<sub>2</sub> (mmHg) | 39.6±3.1 | 41.8±4.3 | 0.38 |
| PH | 7.43±0.05 | 7.42±0.01 | 0.75 |

Note: HR=heart rates; SBP=systolic blood pressure; DBP=diastolic blood pressure; RR=respiratory rates; SpO<sub>2</sub>=pulse oxygen saturation; PaO<sub>2</sub>=partial pressure of oxygen, and PaCO<sub>2</sub>=partial pressure of carbon dioxide.

### UWS dataset

Forty-nine UWS participants (age = 48.88 ± 15.62 years, 13 women; aetiology = 25 stroke, 18 traumatic brain injury, 6 anoxia) with Glasgow Coma Scale (GCS) score (Teasdale and Jennett, 1974) from 3 to 10 and Coma Recovery Scale-Revisited (CRSR) score (Giacino et al., 2004) from 1 to 8 were included in this study. EEG data were acquired for at least 5 minutes using a 256-channel system (GES 300, Electrical Geodesics, Inc., USA) and a 256-channel electrode cap (HCGSN 257-channel net cap, Electrical Geodesics, Inc. USA). EEG signals were acquired at a sampling rate of 1000 Hz and referenced to Cz. The impedance of all electrodes was kept below 20 KΩ.

### ALS dataset

A total of twelve ALS patients (age = 57.88 ± 13.24 years, 7 men, 1 woman, 4 n.a.) with ALSFRS-R score from 3 to 40 (min = 0, max = 48; Cedarbaum et al., 1999), as well as a single female ALS patient (age = 52 years) suffering from LIS (ALSFRS-R = 1) participated in the study. EEG data were acquired for 5 minutes (eyes-open) using 121 active electrodes at a sampling frequency of 500 Hz (Brain Products GmbH, Germany). The placement of the electrodes followed the international 5-10 system, reference to the left mastoid. For a further description of the acquisition procedure, see (Fomina et al., 2017; Hohmann et al., 2018, 2016). The same protocol was undertaken by twenty-three healthy participants in awake condition.

### Ethics Statement

All participants (or their legal guardians) provided informed written consent before participation. This research was approved by the respective Universities/Hospitals depending on the origin of the dataset (Western University Health Science Research Ethics Board for the sleep dataset, Huashan Hospital, Fudan University for the anaesthesia and UWS datasets, and Max Planck Society’s Ethics Committee for the ALS dataset). This study was conducted in accordance with the Declaration of Helsinki guidelines.

### Data/code availability statement

All data are available and will be put into repository. All software and codes are freely available.

### Pre-processing

Given the variety of datasets from different equipment and conditions used in the present study, the specific pre-processing procedure was carried out for each dataset depending on the particular requirements of the data. We took special care in removing muscular and ocular artefacts in the case of anaesthesia, UWS and ALS datasets. For that purpose, EEG signals were bandpass filtered between 0.5 and 40 Hz using a finite impulse response (FIR) filter. Then, independent component analysis (ICA) was applied to remove components from the muscular and ocular artefacts. On the other hand, due to the length of the EEG recordings (polysomnography during a normal sleep), epochs labelled as noise epochs by a registered technologist were completely removed from the data. In the remaining epochs, FIR filter between 0.5 and 40 Hz was applied to the data. All recordings were re-referenced to the average activity. Further details of the pre-processing are explained in Supplementary Material.

### Temporal analysis

After pre-processing, the ACW was computed for each of the participants from the four datasets. For that purpose, custom scripts were developed to compute the ACW by measuring the full-width-half-maximum of the temporal autocorrelation function of each electrode, following the description provided by Honey et al 2012 (see also Fig 1 on the left). Autocorrelation was calculated using windows of 20 second-length with and overlap of 50%. The lag was set to 0.5 seconds since we observed in a previous study that the ACW values agreed for different lag values (ranged from 0.1 to 1 seconds) (Wolff et al., 2019). The full-width-half-maximum of the main lobe of each the autocorrelation functions was then computed for each epoch. ACW was estimated as the average of all the epochs for each electrode and condition. In order to reduce the number of comparisons and to minimize type I errors, a grand average across electrodes was performed. ACW values represent the extent of the periodicity of the EEG signal, whereby longer ACWs can be interpreted as greater stability of the frequencies over time (Fig 1 on the left). The length of the ACW can be seen, therefore, as an index that summarizes the degree of regularity of a signal, with longer ACW associated with more regular EEG oscillations. On the contrary, considering the extreme case, the autocorrelation of a white noise signal will have a peak in the origin, whereby the ACW, in this case, would be zero.

**Fig 1.**
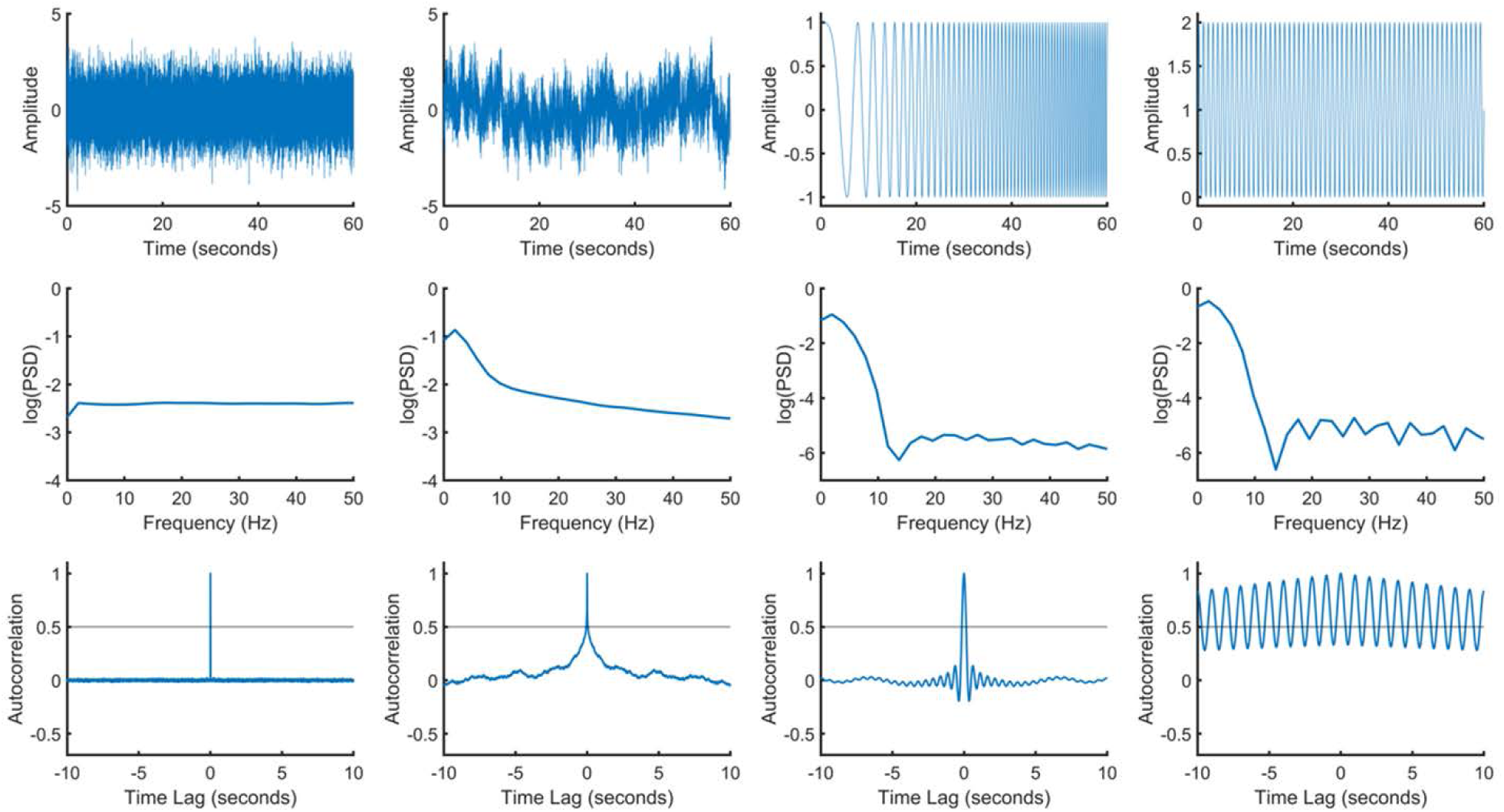
Simulation examples showing the dissociation between PSD and ACW. Different signals are shown in the upper row (some of them usually present on the EEG). The second row shows the autocorrelation function for each of the above signals. Autocorrelation window (ACW) is almost zero for pink noise and white noise. The bottom row shows the power spectral density (PSD) of each signal. The sine wave and the chirp signal show a very similar power-law exponent (PLE). The procedure for obtaining the ACW and the PLE is shown on the left.

### Spectral analysis

To estimate the PSD of the EEG data, Welch’s method was computed (Welch, 1967). This method requires a split of the EEG time series into overlapped segments of length *L*. For our analyses, *L* were set to 3 times the sampling rate, (e.g., 3 seconds), with an overlap of 50%. Then, segments were smoothed using a Hamming window. Fast Fourier Transform (FFT) was applied in an epoch-based way to obtain the modified periodogram. Finally, the PSD was estimated by averaging all the periodograms. This allows us to obtain an adequate resolution (two data samples per Hz) with an assumable increase of the computational cost. PSD values represent the power of oscillatory neuronal activity across the frequency spectrum.

Once the PSD was computed, PLE was obtained using in-house Matlab scripts. For that purpose, PSD representation was log-log transformed in both the frequency and the power spectrum range. Then, the slope of the PSD was estimated computing linear least squares regression. Finally, the PLE of each was defined as the absolute value of such slope (see Fig 1 on the left). The averaged PLE across epochs and channels was used for further analyses. PLE values represent the extent of broadband arrhythmic neuronal activity in the EEG. Thereby, lower PLE values, i.e. more flatness in the PSD function, is associated with a more arrhythmic activity. The extreme is again a white noise signal, with a completely flat PLE.

It is worth noting that the PLE complements the PSD analysis by identifying differences in the temporal structure of the spectrum power. While the PSD shows the differences of the power spectrum in terms of the absolute power at particular frequencies, the PLE instead highlights the specific relationship in power between slow and fast frequencies, showing how their balance is altered in certain states, e.g. in anaesthesia (Zhang *et al*. 2018). For this reason, the increase in power of slower frequencies is not always and necessarily associated with higher negative slope of the PSD (i.e., higher PLE) and vice versa. For example, a PSD that shows high power in slower frequencies may be associated with low PLE (flat slope) in case of an increased power also in faster frequencies. On the other hand, a PSD that shows low power in slower frequencies may be associated with higher PLE in case of an excessive decrement in the faster frequencies.

### Statistical analysis

Statistical analysis was done with Matlab ‘Statistics and Machine Learning’ Toolbox (version 2017b). For parametric data, paired two-tailed *t*-tests were used for within-group comparisons, whereas independent two-tailed *t*-tests were used for between-group comparisons. In the case when parametric assumptions were not met, Mann-Whitney *U*-tests and Wilcoxon signed rank tests were used for within and between-group comparisons, respectively. In the particular case of the sleep dataset, comparison among 5 different conditions (i.e. awake, N1, N2, N3 and REM) were assessed. Since the data did not meet parametric assumptions, the Friedman test was applied. It is important to note that some of our comparisons require more than a conventional superiority test. In particular, we sometimes want to check the equivalence between two different distributions (i.e. H1), which is the opposite of the conventional goal. In this case, the null hypothesis (H0) would be the contrary (both distributions are different) and equivalence or non-inferiority testing are required (Walker and Nowacki, 2011). In this study, we select the more restrictive (equivalence testing), which is tantamount to applying two traditional one-side tests (Walker and Nowacki, 2011). Equivalence margin was set to 50% of the PLE and the ACW differences between awake and N1 in the sleep data. This value was chosen following the most restrictive recommendations of the FDA in mortality studies (Walker and Nowacki, 2011). This procedure allows us to minimize type I errors. Finally, for correlations, Spearman’s rho test was used since we do not have a priori hypothesis about the type of relationship between the variables (i.e. linear or non-linear relationship).

Finally, aimed at controlling for possible bias due to the variety of dataset used, we performed a frequency-to-frequency analysis to reveal spectral bands with significant differences between groups (see Supplementary Material for details).

## Results

### The subtle differences between power spectral density and autocorrelation window

The relationship between autocorrelation function and power spectral density is well-known. In fact, Blackman-Tukey approach (Blackman and Tukey, 1958), which is based on the Wiener-Khinchin theorem (Kay, 1988), states that the Fourier transform of the autocorrelation function of a time series is equivalent to the power spectral density (PSD) of such time series. However, the association between ACW and PLE is far from being obvious. The ACW (defined as the full-width-at-half-maximum of the autocorrelation function) and the PLE have been previously used in several neuroimage studies due to its ability to identify subtle differences between time series that the PSD is not able to recognize depending on the spectral resolution of it (Honey et al., 2012; Walden and Zhuang, 2019; Wolff et al., 2019). Nonetheless, both measures have not directly compared before, remaining unclear their similarities and differences. To illustrate their behaviour and to better understand the main findings of this study, a number of simulations were conducted. In particular, four different 1-minute length time series were synthetically generated: pink noise, sinusoidal wave of 10 Hz, white noise and up-chirp signal. These signals were chosen for their characteristics (differences and similarities in PLE and ACW that help illustrate their relationship) and for being present, to a greater or lesser extent, in the EEG. All of them were generated with 500 points per second (simulating a sampling rate of 500 Hz). The time series of the mentioned signals, along with PSDs and the autocorrelation functions of such signals are shown in Fig 1.

In view of the figures, we can claim that each signal contributes differently to the total PLE and ACW of the EEG. For example, white noise ideally has an ACW close to zero and flat frequency response, therefore, its contribution to both measures is quite limited. On the other hand, pink noise, whose contribution to the spectral structure of the EEG is significant, shows an autocorrelation window close to zero, but a relevant contribution to PLE. Sine-like waves (sine wave at 10 Hz and chirp) usually have lower power than pink noise on the EEG, but their contribution to signal pre-periodicity (and therefore to ACW) is not negligible.Apart from their different contribution of the signals to the total PLE and ACW, the simulations reveal a dissociation between both measures. On the one hand, we can see that white and pink noise signals differ in the time and in the spectral domain, showing a higher contribution of low frequencies in the pink noise signal. These differences are reflected in the autocorrelation function, showing larger ACW for pink noise (0.077) as compared with white noise (0.002). In fact, as previously mentioned, the ACW of a perfect white noise is zero. It is noteworthy that, due to the random nature of these two signals, the ACW was computed as the average of 100 surrogate data. This example reflects the influence of low frequencies in the ACW. On the other hand, the up-chirp signal and a simple sinusoidal wave were also analysed. In this case, the signal periodicity of the sinusoidal wave is higher than the chirp. However, as Fig 1 shows, the PSD of both signals are very similar showing, in both of them, an important influence of low frequencies. Despite the high degree of similarity in both PSDs, the periodicity of the signal is directly reflected in the autocorrelation functions, showing more than double ACW value for the sinusoidal wave (0.666) than the up-chirp (0.306). These simulations reflect the influence of the signal periodicity in the ACW, which makes it useful for measuring subtle differences in the signals in particular cases. At the same time, this example shows the theoretical different contribution of the signals as well as the dissociation between ACW and PLE.

### Intrinsic neural timescales in sleep

First, the PSD for all the sleep stages were estimated (see S1 Fig in the Supplementary Material for PSD representations) and visually inspected. Then, the ACW and the PLE were computed. Our results showed a significant increased length of the ACW values as the sleep stages become deeper (Friedman test, *χ*^2^(4) = 79.63, *p* <0.001). Interestingly, the ACW in wake was significantly lower than in N2 (Wilcoxon signed-rank test, *z* = -4.29, *p* <0.001) and in N3 (Wilcoxon signed-rank, *z* = -4.28, *p* <0.001). The ACW in REM was also significantly lower than N3 (Wilcoxon signed-rank test, *z* = -4.26, *p* <0.001), but not significantly different from N2 (Wilcoxon signed-rank test, *z* = 1.17, *p* >0.05) (see also Khalighi et al. 2013) (see Fig 2a). Analogous changes were observed in the PLE (Friedman test, *χ*^2^ = 87.1, *p* <0.001). Importantly, as with the ACW, we could clearly observe increase of the PLE from the awake state over N1 (Wilcoxon signed-rank test, *z* = -4.14, *p* <0.001), N2 (Wilcoxon signed-rank test *z* = -4.29, *p* <0.001), N3 (Wilcoxon signed-rank test, *z* = - 4.29, *p* <0.001) and REM (Wilcoxon signed-rank test, *z* = 1.24, *p* <0.001), with more noticeable differences with N2 (Fig 2b). Finally, in both the ACW and the PLE, global changes were observed in the topographical maps (Fig 2c and 2d), which is supported by statistical differences found both for the ACW and for the PLE (see S5a Fig and S5b in the Supplementary Material). Note that for Figure 2 and the subsequent figures of each dataset, graphics and topoplot colormaps were adjusted to permit a fair comparison between healthy controls and pathological/abnormal groups.

**Fig 2.**
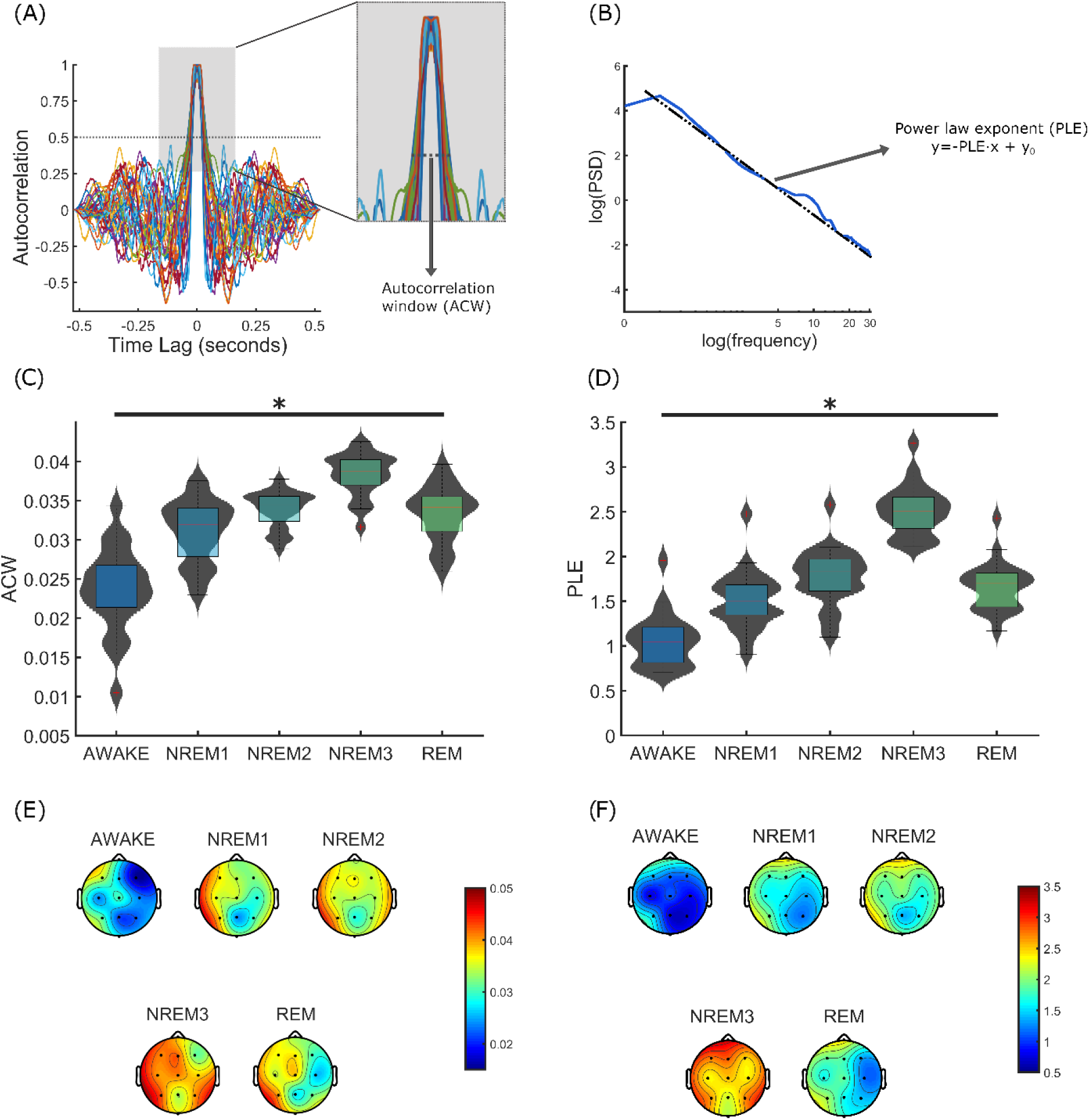
ACW and PLE assessment in the sleep dataset. Illustrative explanation of the ACW (**A**) and the PLE (**B**) computation is shown. The ACW distribution (**C**) and the PLE distribution (**D**) are depicted using violin plots and boxplots for each of the sleep stages. A significant increase for both the ACW and the PLE is observed for deeper sleep stages (Friedman test). Topographical maps for the ACW (**E**) and the PLE (**F**) are also represented for all the sleep stages, indicating global changes which are supported by statistical differences found both for the ACW and for the PLE (see S6a Fig and S6b in the Supplementary Material).

### Intrinsic neural timescales in anaesthesia

The participants receiving sevoflurane (OAA/S = 1) showed the ACW and the PLE values that doubled those shown by the same participants in the awake condition. This is comparable with the results found for N3 in the sleep dataset. Wilcoxon signed-rank tests showed statistically significant differences between awake and sevoflurane conditions in both the ACW (*p* = 0.0125, *z* = 2.50) and the PLE (*p* = 0.0051, *z* = 2.80) for the mean distribution of all the electrodes, demonstrating their abnormally high values in the anaesthetic states as compared to the same participants’ awake state (Fig 3a and 3b).

**Fig 3.**
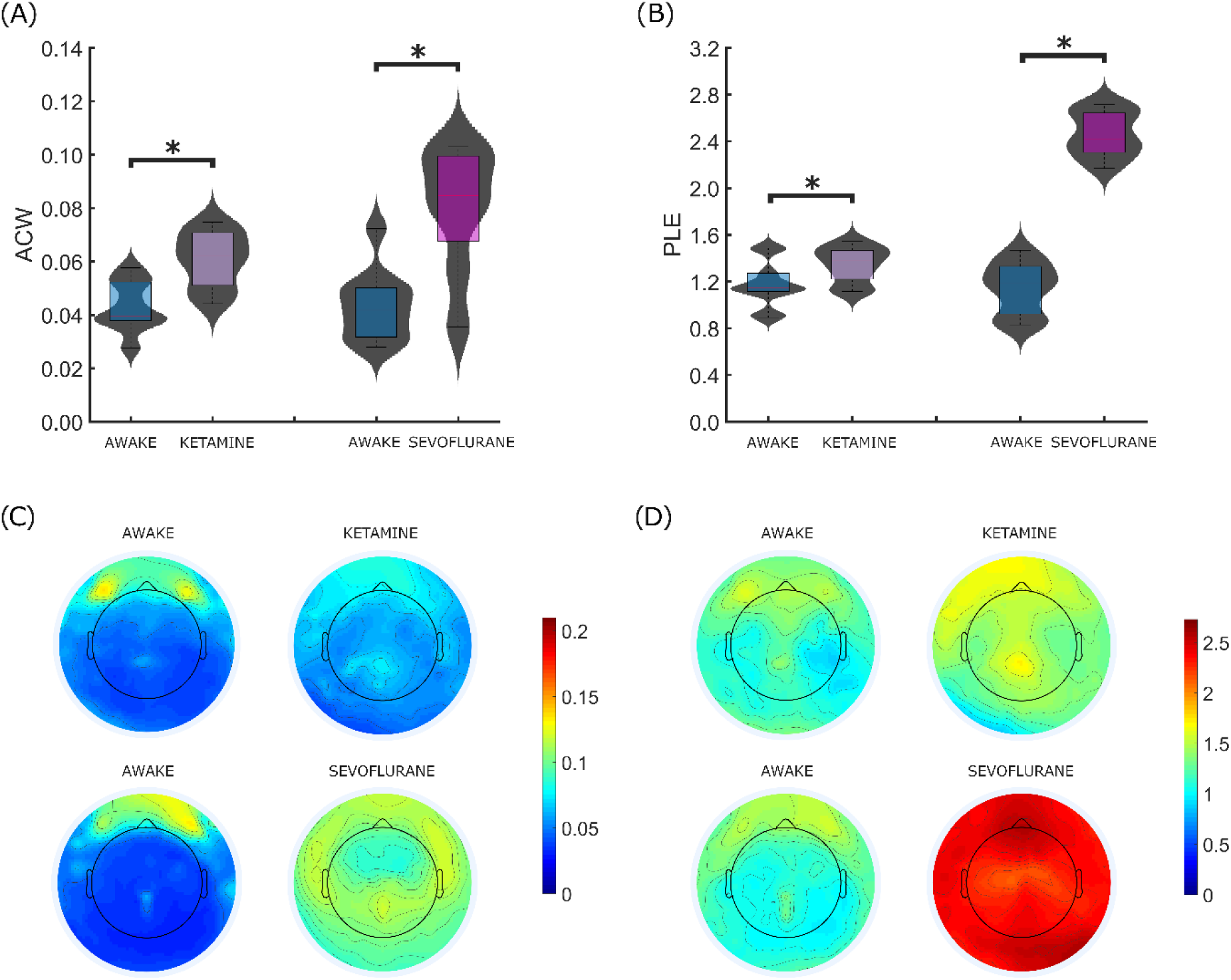
ACW and PLE assessment in the anaesthesia dataset. The ACW distribution (**A**) and the PLE distribution (**B**) are depicted using violin plots and boxplots for anaesthesia and non-anaesthesia conditions. Non-inferiority testing showed non-equivalence between ketamine and awake conditions both for the ACW and the PLE. In addition, a significant increase for both the ACW and the PLE is observed for sevoflurane as compared to awake (Mann-Whitney U-test). Topographical maps for the ACW (**C**) and the PLE (**D**) are also represented for awake and anaesthesia conditions, showing statistical differences in all the scalp in sevoflurane conditions (see S7 Fig in the Supplementary Material). On the contrary, the effects related to ketamine condition are more focused on the parieto-occipital brain region (see S7 Fig in the Supplementary Material).

Similar patterns were found for ketamine condition (OAA/S = 1), both for the ACW and for the same participants’ awake state. However, the increases in the ACW and the PLE as compared to the awake state were much less noticeable than in the sevoflurane group. In this case, no significant differences were found for the ACW (Wilcoxon signed-rank test, *z* = 1.27, *p* >0.01) and PLE (Wilcoxon signed-rank test, *z* = 1.24, *p* = 0.01). Although the ACW and the PLE in the ketamine condition were not significantly different from the awake state, non-inferiority testing showed non-equivalence between ketamine and awake conditions (see Supplementary Material for further details). In other words, ketamine changes enough the ACW and PLE values enough to be considered relevant as compared to placebo conditions. However, these changes cannot be considered significant as compared to the awake condition when the mean of all the electrodes are compared.

In other to further characterize these changes, the same procedure was applied but using the joint distribution of all electrodes (not only the mean). In this case, significant differences were found both for the ACW (Wilcoxon signed-rank test, *z* = 18.03, *p* <0.001) and the PLE (Wilcoxon signed-rank test, *z* = 25.21, *p* <0.001) as compared to awake condition. These differences between sevoflurane and ketamine effects agree with the visual inspection of the PSD, where sevoflurane showed an overall steep decay of the PSD compared to the awake state, with higher PSD values in the slow frequencies (1-8 Hz), a flattening of the alpha peak and a large PSD slope in the higher frequencies (20-40 Hz).

On the contrary, ketamine showed a slight flattening of the PSD, with a slowdown and a shift of the alpha peak towards lower frequencies (see S2 Fig in the Supplementary Material). Finally, as in sleep, the topographical maps showed global effects rather than regionally specific changes for sevoflurane (Fig 3c and 3d), showing statistical differences across the entire scalp (see S6 Fig in the Supplementary Material). On the contrary, the effects related to ketamine condition are more focused on the parieto-occipital brain region (see S6 Fig in the Supplementary Material).

### Intrinsic neural timescales in UWS

Similar to sleep and sevoflurane anaesthesia, the ACW was significantly longer in UWS as compared to the healthy controls (Mann-Whitney *U*-test, *U* = 216, *p* < 0.001) (Fig 4a). Analogously, the PLE was also significantly higher in UWS as compared to healthy controls (Mann-Whitney *U*-test, *U* = 205, *p* = 0.0013) (Fig 4b). Interestingly, Chi-square test showed a weak but significant association between the ACW and symptom severity on the CRSR scale (*χ*^2^(48) = 168.04, *p* = 0.041). In particular, higher ACW values were associated with lower CRSR scores (Pearson’s rho test, *r* = -0,1747). Thus, suggesting that alterations to the intrinsic neural timescale are associated with the severity of disruptions to consciousness. These results are in line with the visual inspection of the PSD, which showed an overall steeper decay compared to healthy participants, with more power in the slow frequencies and a complete flattening of the alpha peak (see S3 Fig in the Supplementary Material). Finally, as in sleep, the topographical maps for AWC and PLE showed a global effect without any specific regional changes (Fig 4c and 4d), which is supported by statistical differences found in most of the electrodes both for ACW and for PLE (see Fig S7 in the Supplementary Material). Material).

**Fig 4.**
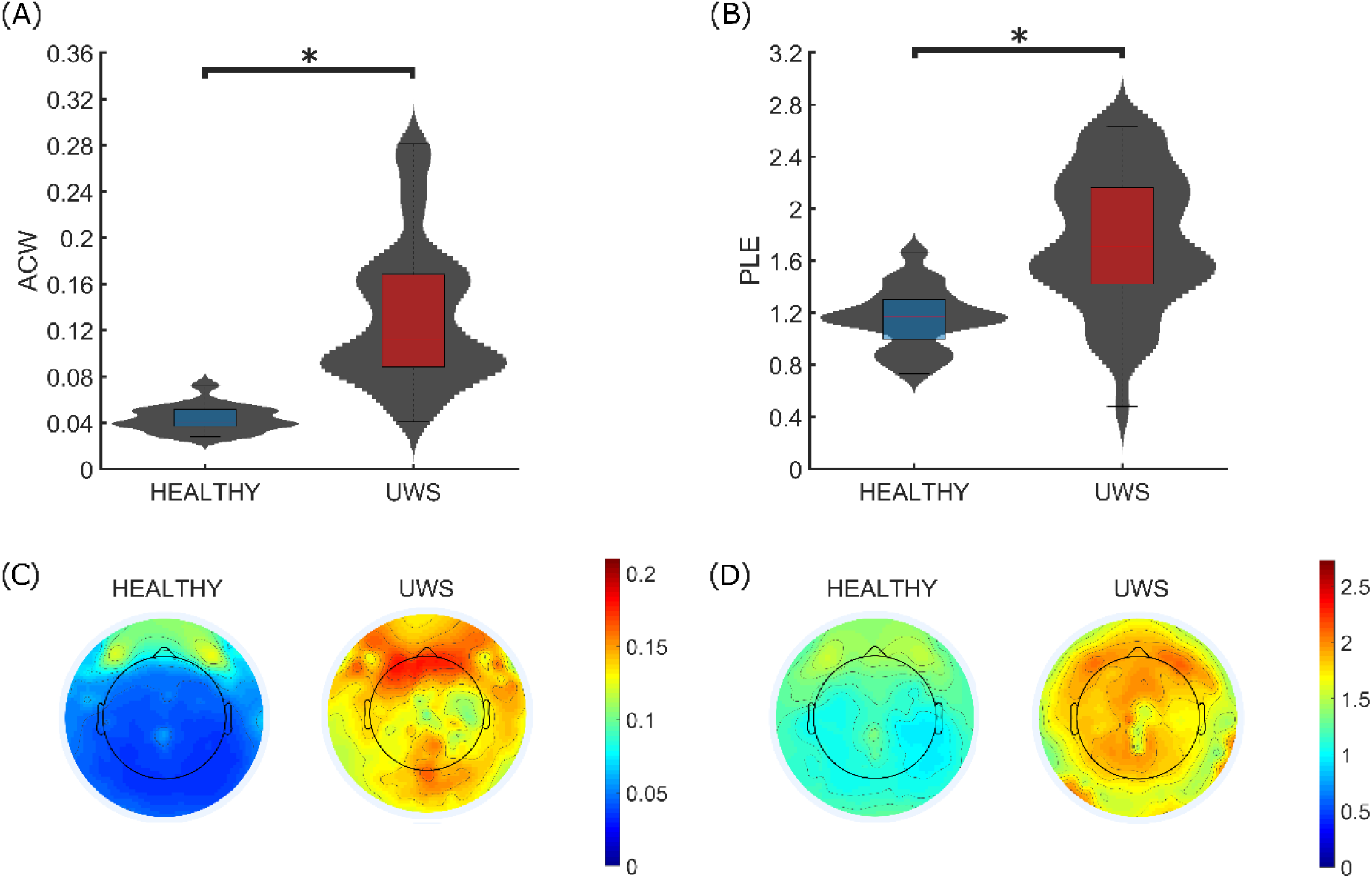
ACW and PLE assessment in the UWS dataset. The ACW distribution (**A**) and the PLE distribution (**B**) are depicted using violin plots and boxplots for UWS participants and healthy controls. A clear increase for both the ACW and the PLE is observed for UWS participants. Topographical maps for the ACW (**C**) and the PLE (**D**) are also represented for healthy controls and UWS participants, indicating regionally no specific effects supported by statistical differences found in most of the electrodes both for the ACW and for the PLE (see S8 Fig in the Supplementary Material).

### Intrinsic neural timescales in ALS with and without locked-in syndrome (LIS)

We also investigated a unique group of participants suffering from ALS with and without LIS. Despite their motor impairment, no significant differences in the ACW (Mann-Whitney *U*-test, *U*= 487, *p* >0.01) or the PLE (Mann-Whitney *U*-test, *U*= 162, *p* >0.01) were found between ALS without LIS and healthy controls (Fig 5). Importantly, although only one individual with LIS participated in this study, this participant showed ACW and PLE values in the same range as both ALS (without LIS) and healthy controls (Fig 5a and 5b). To statistically assess this, *z*-score normalization was performed over the ACW and PLE values of the LIS participant. In all cases, normalized values showed less than one standard deviation from the ACW of the healthy controls (*z* = -0.7181), the ACW of the ALS participants without LIS (*z* = 0.0802), the PLE of the healthy controls (*z* = -0.2633) and the PLE of the ALS participants without LIS (*z* = 0.4285).

**Fig 5.**
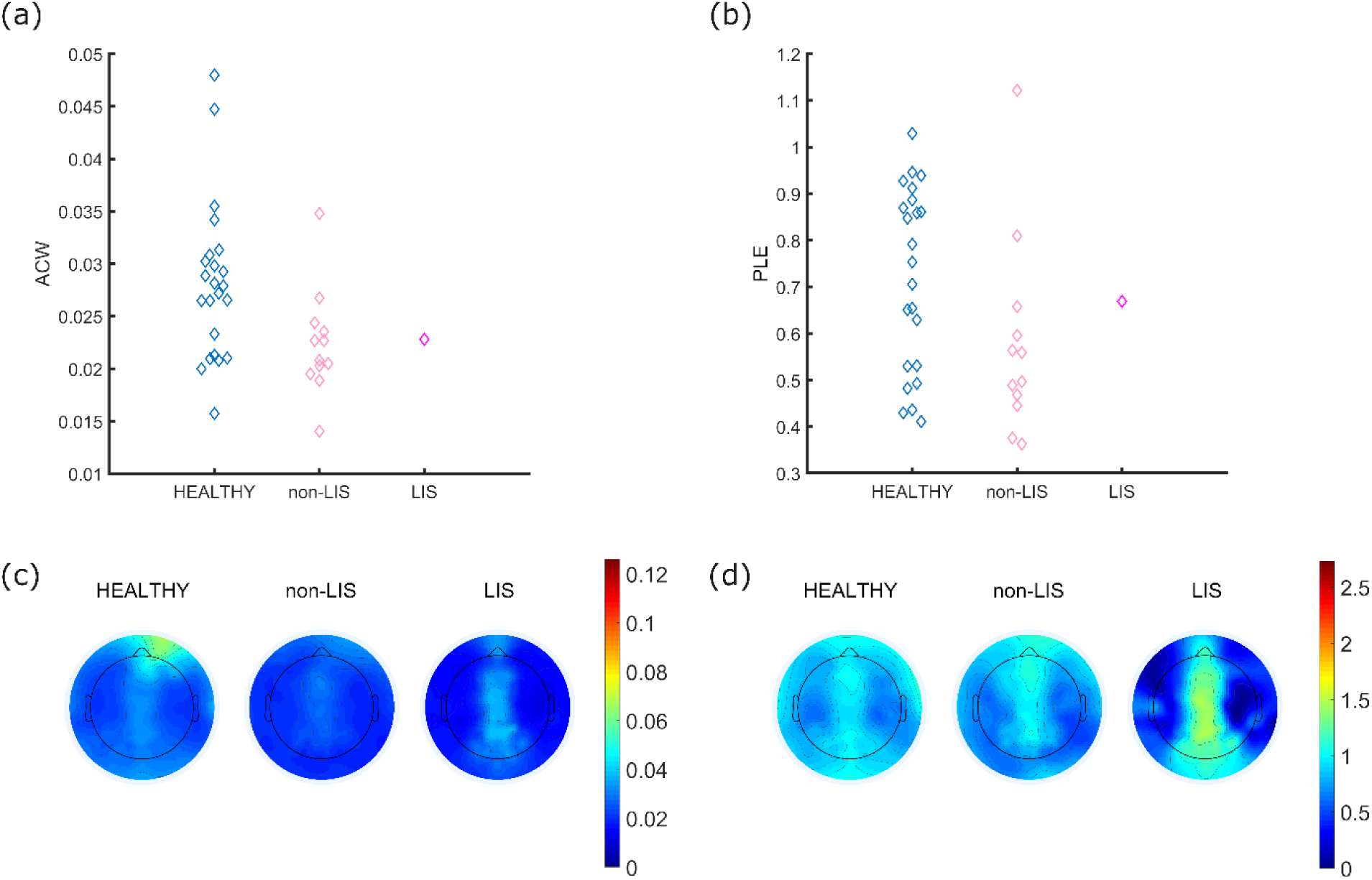
ACW and PLE assessment in the ALS dataset. Due to the number of participants, The ACW distribution (**A**) and the PLE distribution (**B**) are depicted using beeswarm plots for healthy controls, non-LIS participants and the LIS participant. A similar tendency is shown for the three groups for both the ACW and the PLE (non-inferiority testing). Topographical maps for the ACW (**C**) and the PLE (**D**) are also represented for healthy controls and non-LIS participants and the LIS participant, indicating regionally no specific effects.

Consistent with these results, the visual inspection of the PSD, showed a very similar trend for the LIS participant, the ALS participant without LIS and the healthy controls (see S4 Fig in the Supplementary Material). Again, no specific topographical changes were observed (Fig 5c and 5d), which is supported by the regional statistical assessment (see Fig S8 and S9 in the Supplementary Material). Only slight differences were found in the perimeter area of the occipital region. Taken together, these results suggest that the intrinsic neural timescale for ALS either with or without LIS could not be differentiated from healthy controls.

## Discussion

Using resting-state EEG, in a comparative approach, we investigated the relationship between the intrinsic neural timescale of the brain’s spontaneous activity and sensory and/motor information processing through their manifestation in various sensory-or motor-compromised behavioural states. We compared two types of behavioural states: (i) those where sensory information processing is lost while motor information processing seems to remain largely intact, e.g., sleep, anaesthesia, and UWS, and (ii) those where sensory information processing is preserved with only motor information processing being largely deficient, e.g., ALS and LIS.

As predicted, we demonstrated prolonged intrinsic neural timescales, indexed by the ACW in sensory-deficient but motor-preserved behavioural conditions, e.g., sleep, anaesthesia, and UWS. Changes in ACW were accompanied by shifts towards slower frequencies in the PLE and the PSD. In contrast, and as predicted, we did not observe abnormal values in the ACW (or the PLE and the PSD) in motor-deficient but sensory-preserved behavioural conditions like ALS and LIS. Based on converging evidence from these abnormal and normal behavioural states, we conclude that the spontaneous activity’s intrinsic neural timescales are relevant primarily for sensory rather than motor information processing.

### Sensory vs motor information processing

We observed significant modulation (i.e., longer duration) of the ACW in sleep, anaesthesia, and UWS, whereas no change in the ACW was observed in ALS and LIS. These data are well in line with recent findings of dynamic changes in the brain’s spontaneous activity in sleep, anaesthesia, and UWS (Casali et al., 2013; Demertzi et al., 2019; Huang et al., 2018c, 2016, 2014; Piarulli et al., 2016; Sarasso et al., 2015; Siclari et al., 2018, 2017; Sitt et al., 2014; Tagliazucchi et al., 2016, 2013a; Thiery et al., 2018; Zhang et al., 2018). Our results extend these findings to show a prolongation of the brain’s spontaneous intrinsic neural timescales.

Specifically, our results show that the intrinsic neural timescales of the brain’s spontaneous activity are abnormally extended in all conditions examined where sensory information processing is impaired. This included behavioural conditions in different settings, e.g., normal, healthy physiologic (sleep), pharmacologic (anaesthesia), and pathological (UWS). Despite the remarkable differences between these conditions, all these states showed prolongation in their ACW. Thus, the evidence suggests that changes in the brain’s intrinsic neural timescales are related to one specific feature that is shared by all these conditions, i.e., reduced or absent sensory function, independent of their underlying cause. Consequently, we propose that the prolonged ACW in these conditions reflects the loss of the capacity of the brain’s spontaneous activity to process sensory information due to alterations of the intrinsic neural timescale.

In contrast, we did not observe ACW prolongation in conditions such as ALS and LIS where, despite motor deficits, sensory-based interaction with the environment (and consciousness) remain intact. Albeit indirectly inferred through converging evidence using a comparative approach in a collection of abnormal behavioural states, we therefore suggest that the intrinsic neural timescales of the brain’s spontaneous activity are associated with the capacity to support sensory rather than motor information processing.

Our findings complement the observation that the brain’s intrinsic neural timescales are crucially important for the processing of external sensory stimuli during task-evoked activity; this has been described by temporal receptive window (TRW) (Chen et al., 2017, 2015; Hasson et al., 2015) and the temporal receptive field (TRF) (Cavanagh et al., 2016). Given that the magnitude of task-evoked activity is dependent upon an ongoing spontaneous activity (He, 2013; Huang et al., 2017a; Northoff et al., 2011), one would assume that the intrinsic timescale of the brain’s spontaneous activity also shapes sensory processing during task-evoked activity. This possibility, however, remains to be directly investigated.

### Temporal segregation and integration of sensory information

Given that the ACW measures the degree of correlation of neural activity patterns between different time points, accordingly, a short ACW allows for increased temporal precision as it makes it possible to separate different stimuli at distinct time points (Himberger et al., 2018; Murray et al., 2014). That is especially relevant for sensory information processing as high temporal precision is required to distinguish between different sensory-mediated objects and events in the environment in time (i.e., temporal segregation) (Himberger et al., 2018).

External stimuli in all sensory modalities require fast responses as they are brief and change rapidly, which may facilitate behavioural adaptiveness required during healthy, alert wakefulness. This is also reflected in the shorter duration of the intrinsic timescales in sensory cortex in the healthy brain that exhibits short ACW in both rest and task states (Chaudhuri et al., 2015; Gollo et al., 2017, 2015; Hasson et al., 2015; Honey et al., 2012; Stephens et al., 2013). In contrast, increased ACW indicates a stronger correlation across more distant time points. Such temporal autocorrelation is central for temporal integration (e.g., temporal summing and pooling) of different inputs at a particular point in time (Himberger et al., 2018, p. 163).

Our findings indicate that such temporal pooling and summing are abnormal in those behavioural conditions exhibiting impairment in sensory information processing. The abnormal prolongation of the ACW observed here signifies an increased capacity of the brain’s spontaneous activity for temporal summing and pooling. Different sensory inputs at different points are thus lumped and integrated into the same neuronal event. This would have the consequence of impairing temporal precision and segregation of sensory information processing over time such that different sensory stimuli at different time points are no longer distinguishable from one another. That, in turn, may lead to the loss of temporally-specific and -precise sensory-based responsiveness to the external environment that is shared by all three states, e.g., sleep, anaesthesia, and UWS (but not the motor-deficient states like ALS and LIS).

In addition to the ACW, we also measured the PLE and the PSD. Together, as expected (Casali et al., 2013; Colombo et al., 2019; Demertzi et al., 2019; Huang et al., 2017b, 2016, 2014; Lehembre et al., 2012; Schiff et al., 2014; Siclari et al., 2018; Sitt et al., 2014; Tagliazucchi et al., 2016, 2013a; Tagliazucchi and van Someren, 2017; Zhang et al., 2018), our results show decreases in fast frequency power with a shift towards relatively stronger slow frequency power in both the PLE and the PSD in sleep, anaesthesia, and UWS. In contrast, such a shift towards slower frequencies was not observed in ALS and LIS.

Slow frequencies can be characterized by long cycle duration which distinguishes them from the shorter cycle duration of faster frequencies (Buzsáki, 2006). The long cycle duration renders the slow frequencies ideal for integrating or lumping together different stimuli (He and Raichle, 2009; Northoff, 2017, 2014a, 2014b). We therefore tentatively assume that the shift in power towards slower frequencies with their long cycle durations, as measured with PLE and PSD, in sleep, anaesthesia, and UWS, increases the capacity for temporal integration. Due to the concurrent decrease in the power of the faster frequencies, this would decrease temporal precision and segregation of sensory information processing. This would, in turn, result in a lack of sensory-based responsiveness including conditions such as N1-3 sleep, anaesthesia, and UWS. However, to conclusively support these hypotheses, future studies may be needed.

### Limitations

The brain’s intrinsic neural timescale is supposed to exhibit an intricate hierarchy with sensory regions showing shorter ACW and higher-order prefrontal regions revealing longer ACW (Gollo et al., 2017, 2015; Honey et al., 2012; Kiebel et al., 2008; Murray et al., 2014; Stephens et al., 2013). However, our topographical maps did not reveal the specific location of the ACW changes, thus, it was not possible to explore the spatial topography of the intrinsic neural timescale.

We only included participants suffering from *reduced* or *loss* of consciousness. This leaves open the possibility that future studies might assess ACW in participants with so-called ‘extended consciousness’, e.g., during drug-induced psychosis with LSD, psilocybin, mescaline or others (Atasoy et al., 2018; Carhart-Harris, 2018). Moreover, the findings show that the power of faster EEG frequencies is relatively increased in these states while, at the same time, slow frequency power (in absolute terms) is preserved (Atasoy et al., 2018). One would consequently expect shorter duration of ACW (as its length is then driven mainly by the faster than the slower frequencies) and lower PLE (as the increased power in the faster frequencies lowers the PLE) in extended consciousness conditions. This possibility remains to be investigated.

It is important to note that our different groups are characterized by differences other than sensory vs. motor loss, which was the focus of the current investigation. Therefore, we cannot exclude the possibility that the observed differences between anaesthesia/UWS/sleep and ALS/LIS in terms of the ACW, PLE and PSD are due to factors other than the difference between loss of sensory or motor function. However, it is worth mentioning that the converging evidence from these independent groups and conditions suggest that the intrinsic neural timescale varies in a meaningful way, despite the underlying cause of the conditions, and how they present themselves. To the best of our knowledge, the different datasets analysed here do not vary systematically in some other way that might easily explain the pattern of results.

Due to the diversity of participants and conditions in this study, EEG recordings were acquired using different equipment, which involves a different spatial resolution or sampling frequency depending on the dataset. For that reason, different pre-processing procedures were applied according to the necessities of each condition. Although one can consider it a drawback, the fact of obtaining consistent findings in all the databases increase the robustness, and generalizability of our results.

Finally, we did not go into details about the results in REM-sleep where differences in ACW and PLE were less pronounced and more wake-like. This is consistent with the neurophysiology the neurochemistry and cognitive state of REM sleep, which is paradoxically wake-like (Houldin et al., 2019) in the sense that the EEG resembles that of wake (e.g., high frequency, desynchronized, low amplitude EEG), acetylcholine is high, and, as in wake states vivid mentation characterizes dream content (Siegel, 2011). This suggests that indeed, even within healthy and normal diurnal variations in behavioural states, the temporal dynamics, e.g., ACW, PLE, and PSD, of the brain at rest can be modulated in different degrees on a continuum including a variety of different dynamic states (see also (Northoff et al., 2019; Northoff and Tumati, 2019)).

## Conclusions

Taken together, extending recent findings on temporal receptive windows (TRW) during task-evoked activity, we show that the intrinsic neural timescales of the brain’s spontaneous activity are associated with temporal integration (or segregation) of specifically sensory rather than motor information processing. How the intrinsic neural timescales of the brain’s spontaneous activity including how the relationship to sensory information processing stands in relation to the sensory-based TRW during task-evoked activity remains unclear though. Future studies are thus warranted combining both resting and task states during the investigation of intrinsic neural timescales.

