## Supplementary material for "Intrinsic neural timescales related to sensory processing: Evidence from abnormal behavioural states"

### 1. Details of the pre-processing steps

Due to the diverse conditions of the recordings from the four datasets used in the present study, different pre-processing procedures were carried out. In particular, sleep data show the main differences because of the duration of the recordings and the high non-stationarity of the signals compared to the other datasets. While sleep recordings have an all-night duration, the other datasets last only a few minutes. In addition, the different phases of sleep that a person goes through at night are well known, being the stationarity of the EEG signals lower than during awake (Schulz, 2007). These phases, or sleep stages, have different characteristics in terms of amplitude and frequency of the EEG signal (Schulz, 2007), so splitting into sleep stages and characterize them individually is imperative. The other main difference between datasets is the number of electrodes. Since sleep recordings come from polysomnography (PSG), the number of electrodes is lower than the other datasets. This prevents applying methods based on signal decomposition, such as independent component analysis (ICA). Since the same ICA components are obtained as the number of input channels (Delorme and Makeig, 2004), a large number of channels is necessary to accurately determine the components coming from eye movement and muscle noise.

All the pre-processing was applied using in-house MATLAB (The MathWorks, 2017b) scripts and the EEGLAB toolbox.

#### Pre-processing of sleep recordings

All-nigh sleep EEG recordings from and common PSG were scored into the different sleep stages in epochs of 30 second using RemLogic analysis software (Natus) following the procedure described in (Fang et al., 2017b). During the acquisition. Signals were re-referenced to the contralateral mastoid derivations. Next, data were resampled to 250 Hz to reduce the computational load (well above the Nyquist frequency) and re-referenced to the average activity. After removing noise epochs, signals were filtered using a band-pass finite impulse response (FIR) filter between 0.5 and 40 Hz using a Hamming window. Finally, an adaptative epoch rejection protocol using a statistical-based thresholding method was applied: (i) the mean and standard deviation of each channel was computed, (ii) epochs that exceeded mean ± 4 × standard deviation in at least two channels were discarded (Bachiller et al., 2015; Núñez et al., 2017). For all the steps, custom Matlab scripts were used.

#### Pre-processing of anaesthesia, UWS and ALS recordings

Almost identical pre-processing steps were followed in all these three datasets. First, the sampling rate was down-sampled to 250 Hz using EEGLAB's resample function. Second, the continuous data were band-pass filtered from 0.5 to 40 Hz. The unused channels were then removed, i.e. peripheric channels, as well as channels related to ocular or heart movements. Next, *clean_rawdata* EEGLAB plugin was applied to remove flatline channels, low-frequency drifts, noisy channels and short-time bursts from each EEG channel. Recordings were referenced to the average activity. Finally, stationary artefacts, specifically eye movements and muscular noise, were reduced using ICA.

### 2. Non-inferiority testing

#### Justification

A common goal of many studies is to determine whether two or more distributions are different among them, i.e. to check if a distribution is superior to another. In this type of tests, usually known as superiority test, the null hypothesis (what the investigator hopes to disprove, $H_0$) establishes that there is no difference between distributions. On the contrary, the research hypothesis ($H_1$) is that there is a difference between them. These tests tolerate a type I error fixed by the significance level of the test, which is fixed to 0.05 in many cases.

In this study, however, what we want to prove in particular cases is the contrary than in a usual superiority test. As we stated in the introduction, our research hypothesis, $H_1$, is that there are "no ACW changes in ALS", i.e., the two distributions are equivalent. In this context, the traditional comparative is not valid, and an equivalence testing is mandatory to minimize type I errors (Walker and Nowacki, 2011).

### 3. Procedure

Although it is not a usual practice, a *p*-value can be easily reported using any statistical software. In fact, a *p*-value for equivalence can be calculated as the larger *p*-value of two traditional one-sided tests (Walker and Nowacki, 2011; Wellek, 2010). Like in traditional testing, a *p*-value lower than alpha establish that the data meet the research hypothesis, i.e. equivalence between distributions. In this study, two one-side tests using Mann-Whitney *U*-test was applied to the comparison between healthy controls and ALS participants for both the ACW and the PLE.

### 4. PSD figures

Prior to the PLE estimation, the PSD was computed for each participant, epoch and channel of the four datasets. A grand average (average across participants, channels, and epochs) is shown here as a visual assessment of the differences between conditions.

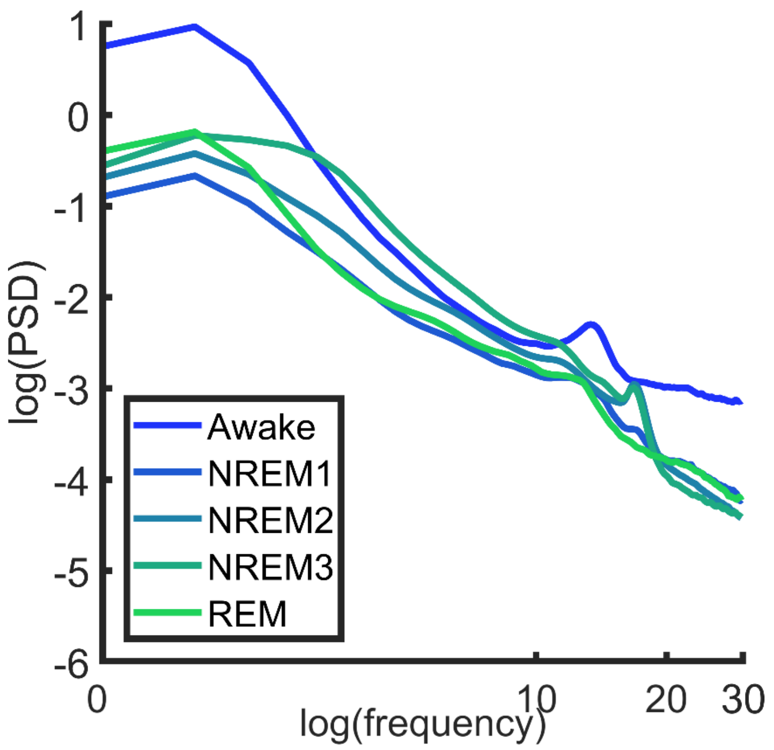


**S1 Fig. Grand average power spectral density (log-log scale) from the sleep dataset.** Related to Figure 2. Mean PSD is represented for awake and all the sleep stages. Notably, despite the higher power in the slow frequencies during the awake state, as shown in the PSD, the structure of the power spectrum changes in N1, N2, N3 and REM, as the relationship in power between the slow and fast frequencies is altered, with a shift towards lower frequencies.

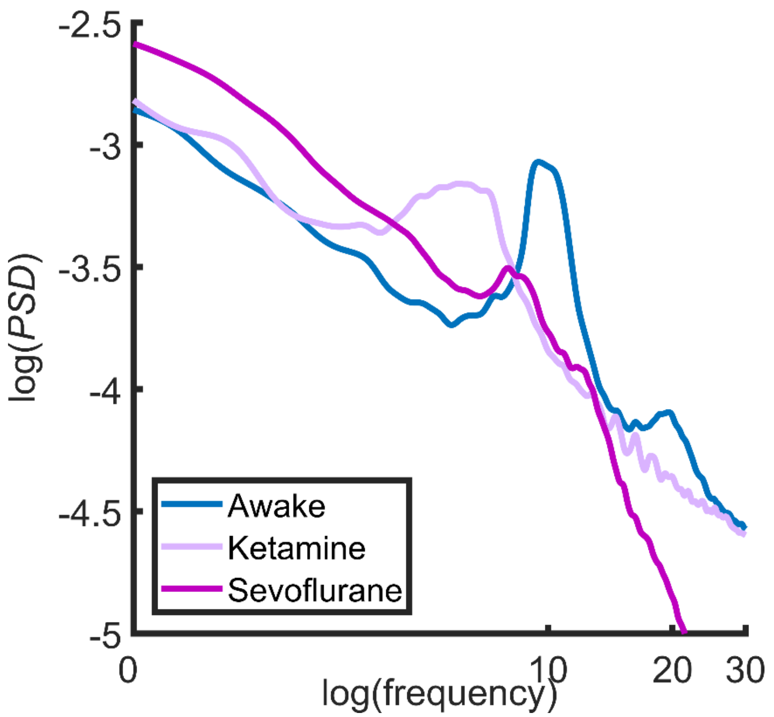


**S2 Fig. Grand average power spectral density (log-log scale) from the anaesthesia dataset.**

Related to Figure 3. Mean PSD is represented for awake and anaesthesia states.

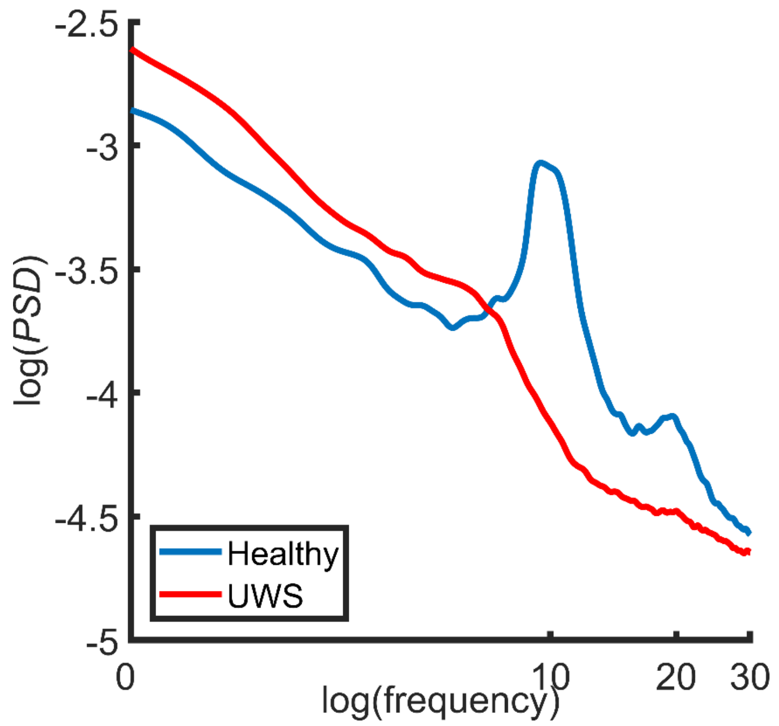

**S3 Fig. Grand average power spectral density (log-log scale) from the UWS dataset.** Related to Figure 4. Mean PSD is represented for healthy controls and unresponsive wakefulness participants.

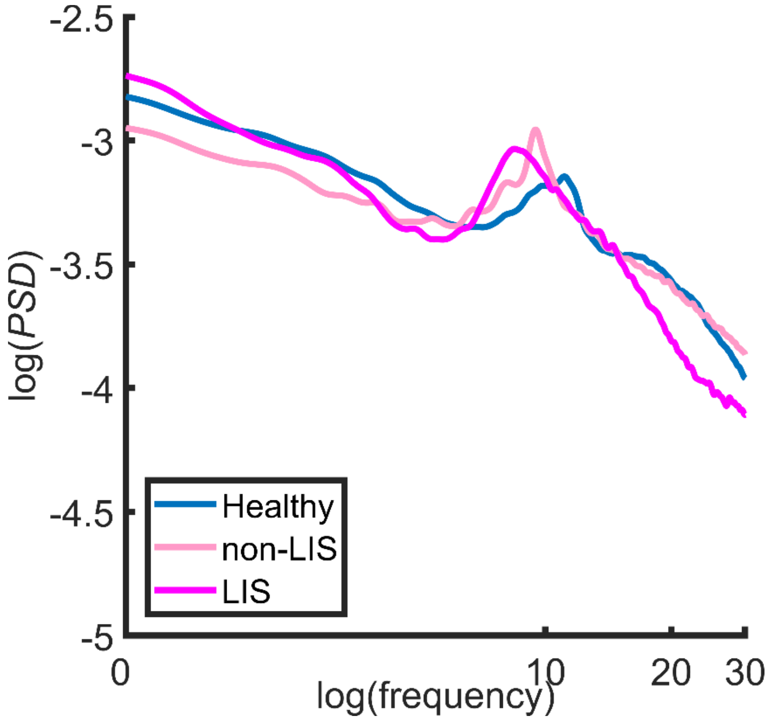


**S4 Fig. Grand average power spectral density (log-log scale) from the ALS dataset.** Related to Figure 5. Mean PSD is represented for healthy controls, non-LIS participants and the LIS participant.

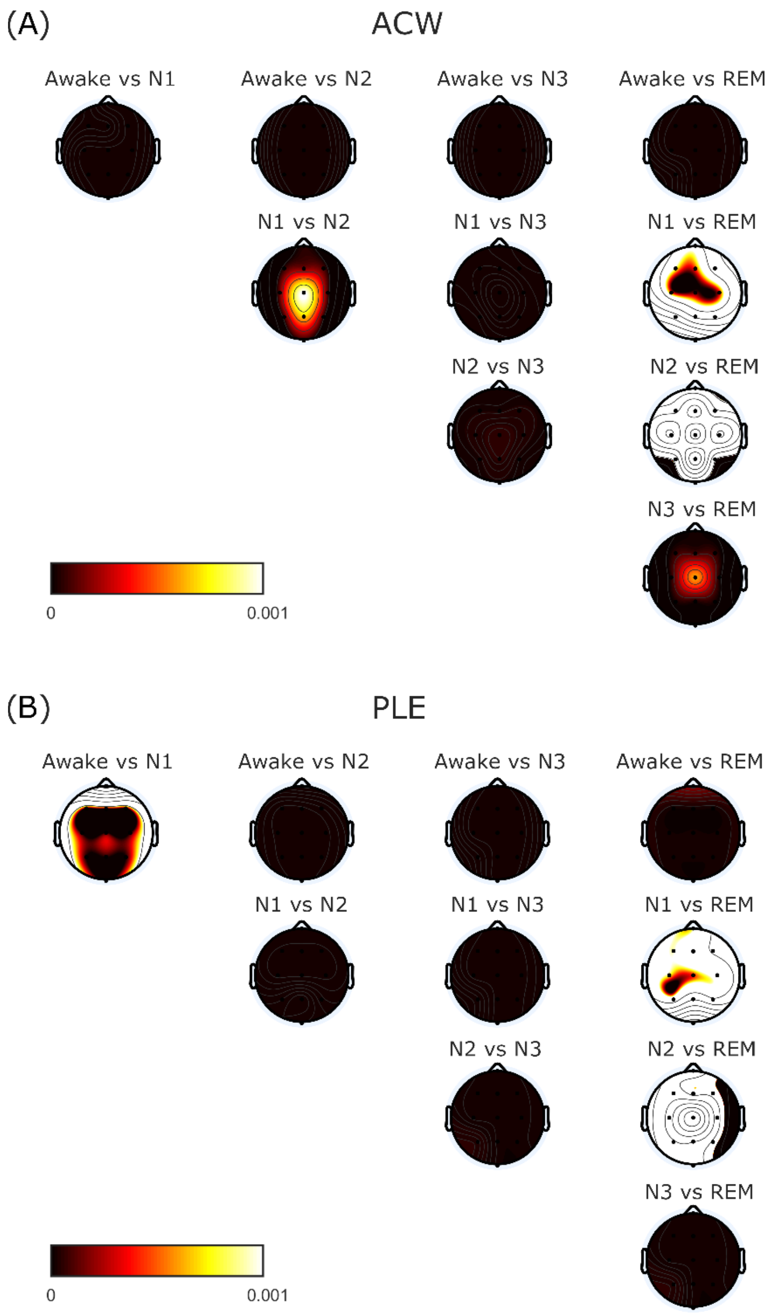


**S5 Fig. Topographical differences for the sleep dataset.** Related to Figure 2. Widespread significant differences were found except for the comparison between N1 vs REM and N2 vs REM both for the ACW (A) and the PLE (B).

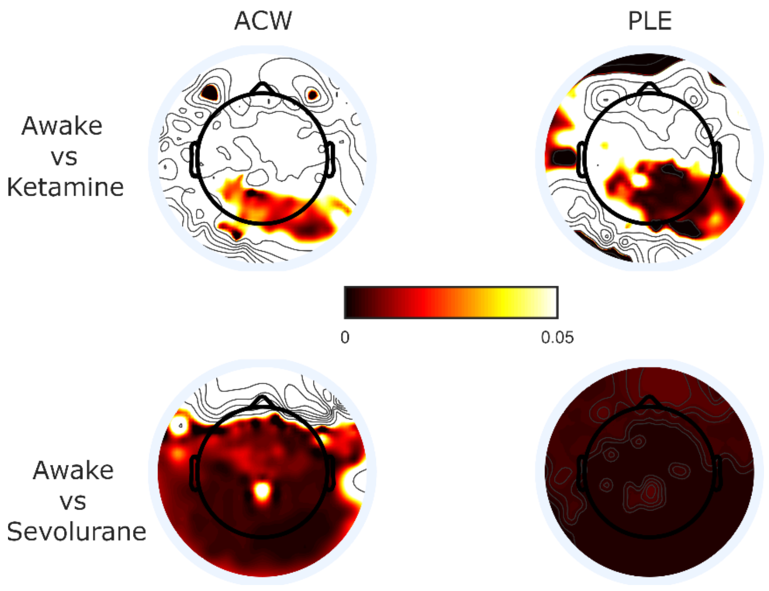


**S6 Fig. Topographical differences for the anaesthesia dataset.** Related to Figure 3. While global significant statistical effects are showed for sevoflurane condition, the effects related to ketamine condition are more focused on the parieto-occipital brain region.

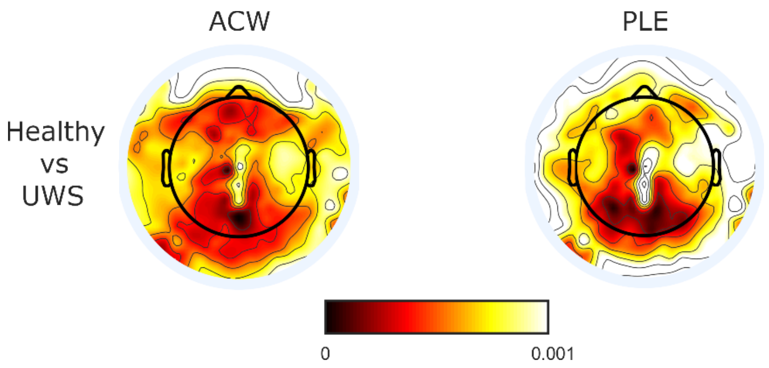


**S7 Fig. Topographical differences for the UWS dataset.** Related to Figure 4. Noticeable statistical differences were found for almost all the electrodes both for the ACW and for the PLE, showing a global effect without any specific regional changes.

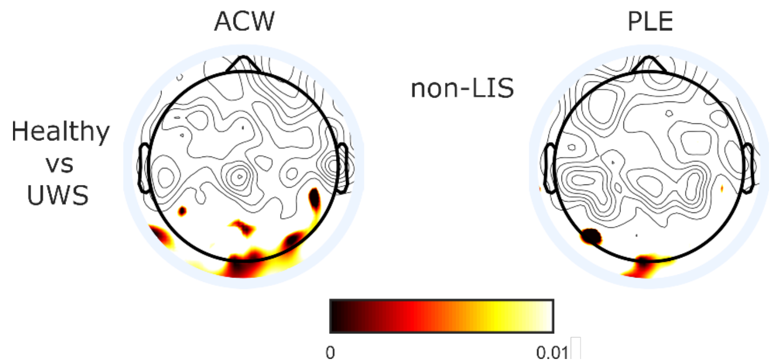


**S8 Fig. Topographical differences for the ALS dataset.** Related to Figure 5. Noticeable statistical differences were found for almost all the electrodes both for the ACW and for the PLE, showing a global effect without any specific regional changes.

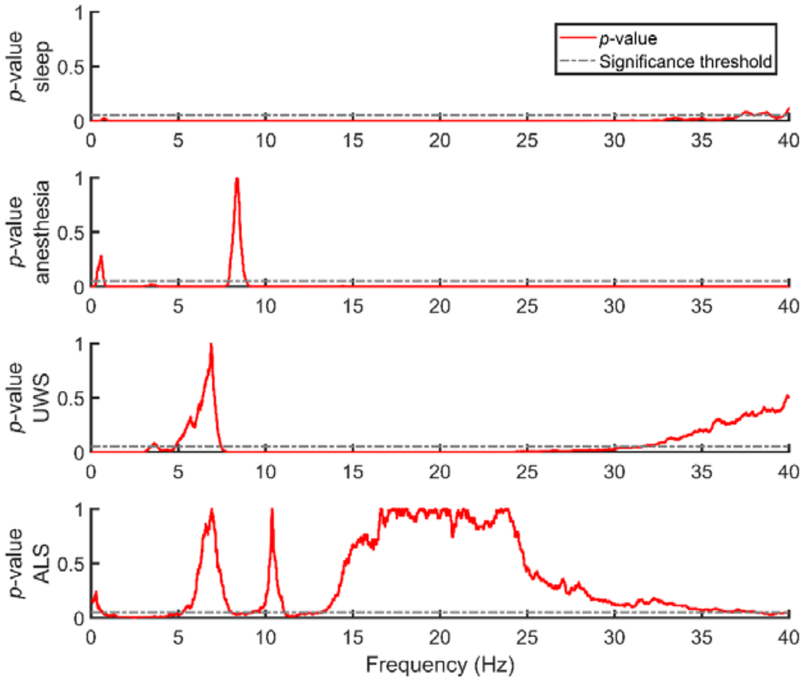


**S9 Fig. Statistical analysis of the significant differences between groups in the spectrum for each of the datasets.** Related to Figures 2, 3, 4, 5. A frequency-by-frequency statistical was performed for each dataset. Frequency bands in which the p-value (red line) is under the threshold (grey line) means statistical differences between groups or conditions. According to the nature of the data, a different test was used: Friedman test (sleep dataset), Kruskal-Wallis test (anaesthesia dataset) and Mann-Whitney test (UWS and ALS datasets).
